# Insulin-like growth factor 1 induces oxidative damage, but does not affect survival in a songbird

**DOI:** 10.1101/2020.03.27.012120

**Authors:** Ádám Z. Lendvai, Zsófia Tóth, Janka Pénzes, Sarah Vogel-Kindgen, Bruno A. Gander, Csongor I. Vágási

## Abstract

Lifespan evolves as a compromise between antagonistic selection forces. Insulin-like growth factor 1 (IGF-1) is a pleiotropic hormone that regulates several life-history traits. High levels of IGF-1 have been linked to increased mortality, partly by causing oxidative stress. However, these effects have no experimental evidence in wild animals. We implanted microspheres loaded with exogenous IGF-1 into bearded reedlings, a common short-lived Eurasian songbird. The treatment elevated plasma IGF-1 levels for at least 24 h. Oxidative damage to lipids significantly increased the day after the manipulation in treated birds, but returned to baseline levels four days post-treatment. The treatment had no effect on survival over 16 months; however, birds with higher pre-treatment (baseline) IGF-1 levels had better survival prospects. These results suggest that, although high IGF-1 levels may induce oxidative damage, natural variation in this hormone’s level may reflect the outcome of individual optimization.

## 1. Introduction

Understanding the regulation of physiological, behavioural, and life-history traits is a central scope of biological research. The ligands of the insulin/insulin-like growth factor 1 (IGF-1) signalling (IIS) pathway stand out as key regulators, because this evolutionarily conserved pathway is present in the whole animal kingdom, and IGF-1 has an antagonistic pleiotropic effect on different fitness components: at high levels, it stimulates growth and reproduction, but impedes self-maintenance processes [1].

Repressed activity of the IIS pathway augment self-maintenance or survival functions resulting in extended lifespan from roundworms and flies to mice and humans [2]. Silencing of the IIS activity extends lifespan in part via increased resistance to oxidative stress ([3]; reviewed by [1,2,4]). However, studies investigating the role of IGF-1 in coordinating fitness and oxidative stress in wild animals are surprisingly scarce [1]. It is still contentious whether a high IGF-1 titre triggers oxidative damage, and this assumption has never been explored in any wild organism [1].

We carried out an experimental study with 40 young bearded reedlings (*Panurus biarmicus*). Our aim was to achieve a sustained increase in plasma IGF-1 levels over a prolonged period (up to four days). We either injected dispersions of IGF-1 loaded microspheres (treated group) or the dispersion medium (control group). We assessed the effect of IGF-1 treatment by measuring oxidative damage to lipids and by monitoring the mortality of individual reedlings in captivity over 16 months.

## 2. Material and methods

### (a) Study species, experimental setup, mortality

Forty juvenile bearded reedlings were caught at Hortobágy-Halastó (N47.6211, E21.0757) and taken into captivity between July 28 and 30, 2017. Birds were initially housed in groups of four individuals in cages measuring 100 × 30 × 50 cm (L × W × H) placed in an outdoor aviary. After at least 10 days of acclimation, birds in each cage were randomly assigned to either IGF-1 or control treatment. Treatments were started in a staggered manner over two weeks to minimize handling times. On the morning of the treatment (day 0), we removed the birds from their cage and took a baseline blood sample within 3 min (time measured from entering the aviary). Then, we injected subcutaneously 100 μL dispersion containing either microspheres loaded with recombinant human IGF-1 (treatment; 2.2 mg microspheres containing 272 ng/mg IGF-1) or only the dispersion medium (control). Dispersion medium consisted of 1.5% (m/m) carboxymethyl cellulose, 5% mannitol, 0.02% and polysorbate 80 in sterile saline solution. Microspheres had been designed to release IGF-1 over several days [5]. Birds were then replaced into their cages. Blood samples were taken after 24 h and 96 h (day 1 and day 4 post-treatment) to assess the short-term physiological effects of the treatment. At three months post-treatment, between November 20 and 22, 2017, all birds were recaptured to take another blood sample for testing long-term repeatability of circulating IGF-1 levels. Birds were then released back into the aviary for additional 13 months (i.e. 16 months in total). Bearded reedlings are short-lived passerines with high juvenile mortality [6]. Therefore, the study period was sufficiently long to detect enough mortality events for statistical analyses. Food and water was provided *ad libitum* and refreshed daily throughout the study [7]. Mortality events were recorded on a daily basis. After 16 months in captivity, on December 8, 2018, all surviving birds (*n* = 12) were released at the site of capture.

### (b) Physiological measurements

Plasma IGF-1 levels were measured by an in-house ELISA assay, as described elsewhere [7]. Plasma malondialdehyde (MDA) concentration reflects the level of peroxidative damage to cell membrane lipids and is a toxic oxidant itself [8]. MDA was measured by high performance liquid chromatography, as detailed elsewhere [8].

### (c) Statistical analyses

All statistical analyses were carried out in R 3.6.2 [9]. We analysed treatment effects on circulating IGF-1 and MDA levels by generalized mixed-effects models (GLMMs) with treatment and sampling time (days 0, 1, and 4) as fixed factors, and individual as the random effect as implemented in package ‘lme4’. We then compared treatment and control groups at each time point by specifying contrasts by the function ‘pairs’ in package ‘emmeans’. Repeatability of IGF-1 level was estimated using the package ‘rptR’. Survival analyses were carried out by Aalen’s regression (function ‘aareg’ in package ‘survival’) that allows for additive effects on the cumulative hazard function. Individuals alive at the end of the study, and one individual that escaped from captivity were right-censored in the models.

## 3. Results

IGF-1 treatment resulted in a transient increase in IGF-1 levels and a corresponding increase in oxidative damage (Fig. 1). IGF-1 levels were similar in the two groups before the manipulation (*p* = 0.8), but higher in the treated group than in controls (*p* < 0.001) on the day after injection of the IGF-1 loaded microspheres. By day 4, this difference between the two groups disappeared (*p* = 0.9). Inter-individual variation in IGF-1 levels remained consistent throughout the study period, resulting in significant repeatability over three months (*R* = 0.30, *p* = 0.029). Similar to IGF-1, MDA levels did not differ between treatment and control groups before the treatment (*p* = 0.2). However, IGF-1-injected birds had higher oxidative damage on day 1 (*p* = 0.021), but this difference disappeared by day 4 (*p* = 0.5, Fig. 1).

**Figure 1.**
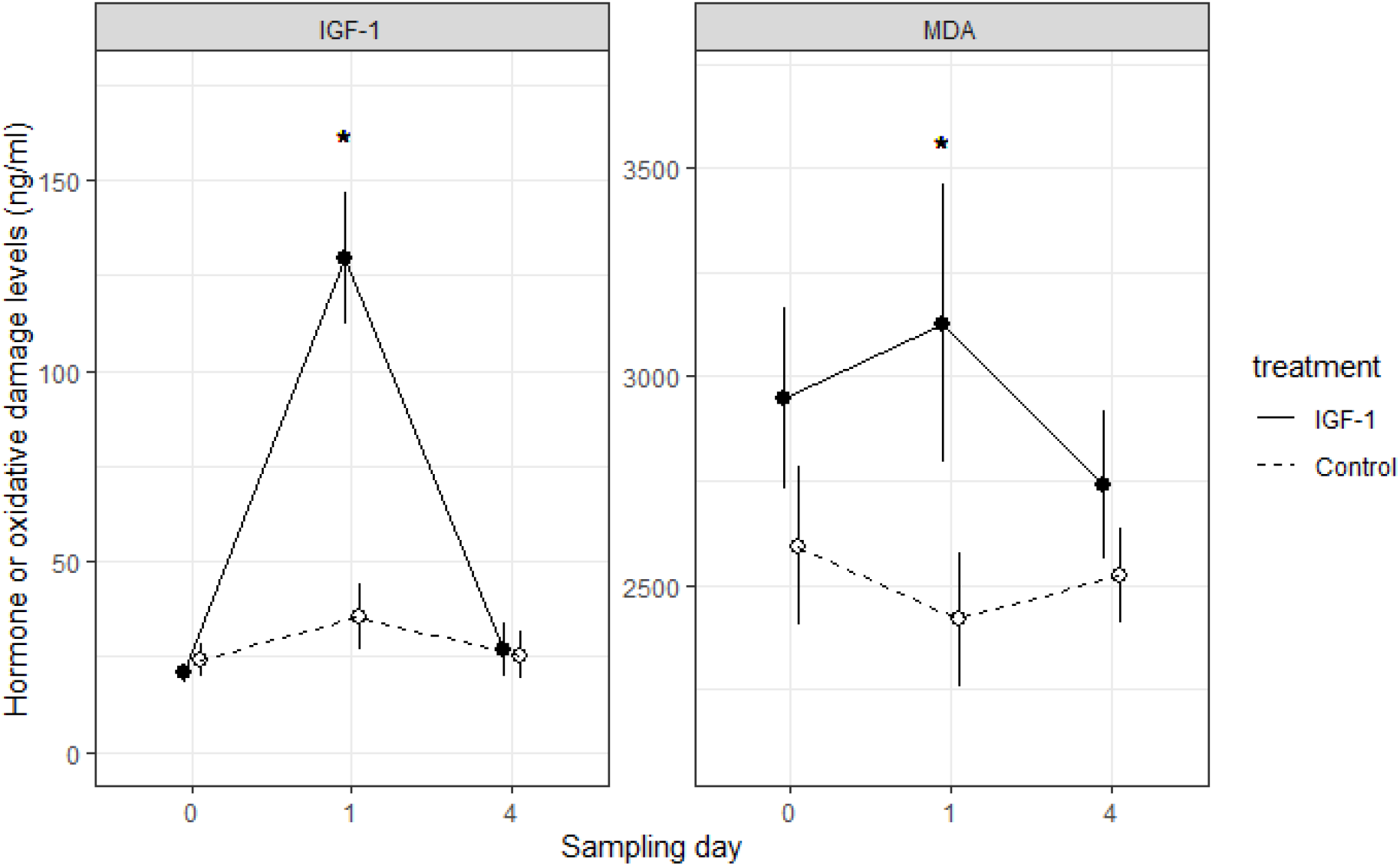
Subcutaneous injection with IGF-1-loaded microspheres resulted in a significant increase in circulating IGF-1 and oxidative damage (MDA) levels measured 24 h later (day 1) in captive bearded reedlings, but these effects disappeared by day 4. Mean ± s.e.m. are shown, asterisks denote significant differences between the treatment and control groups.

Survivorship over 16 months was almost identical in the IGF-1-treated and control groups (*p* = 0.8, Fig. 2); thus, IGF-1 levels on day 1 post-treatment did not affect survivorship. However, birds having higher pre-treatment (day 0) IGF-1 levels were slightly more likely to survive (Table 1). Neither pre-treatment (day 0) nor peak (day 1) MDA levels were associated with survivorship (all *p* > 0.6).

**Figure 2.**
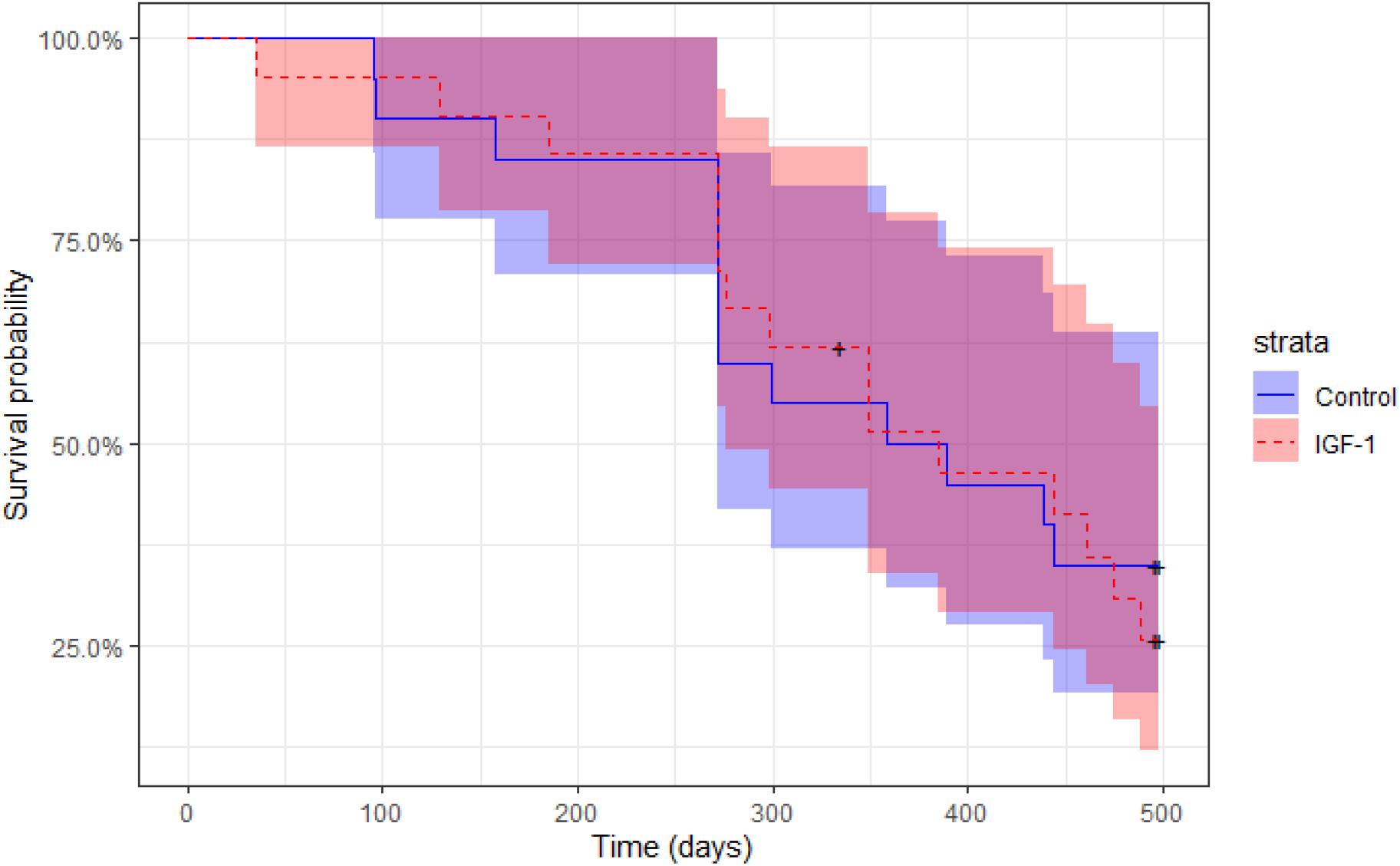
IGF-1 treatment did not affect survivorship in bearded reedlings. The solid lines represent the Kaplan-Meier survival curves, shaded areas denote the corresponding 95% confidence intervals. Cross symbols show censored values.

**Table 1.** Survival model predicts that the likelihood of mortality increases over time, but higher pre-treatment (day 0) IGF-1 levels reduce mortality, while post-treatment IGF-1 (day 1) levels do not affect it.

| fixed effects | estimate ± s.e.m. | *z* | *p* |
| --- | --- | --- | --- |
| baseline hazard | 3.97e-02 ± 0.014 | 2.88 | 0.003 |
| IGF-1 at day 0 | -4.19e-04 ± 0.001 | 1.95 | 0.050 |
| IGF-1 at day 1 | 9.85e-05 ± 0.001 | 0.74 | 0.458 |

## 4. Discussion

IGF-1 is a pleiotropic hormone having antagonistic effects on life-history traits [1], but the adaptive value of variation in its plasma levels remains unknown. Higher IGF-1 titres might be associated with increased mortality in garter snakes, mice, spotted hyenas, and humans, though effect sizes differ between studies and according to the sex and age of individuals [3,10–14]. Although the exact mechanism of such increased mortality remains uncertain, several studies suggested oxidative stress as a mediatory agent ([3]; reviewed by [1,2,4]).

Here, we showed for the first time that the experimental elevation of circulating IGF-1 level caused increased levels of oxidative damage at short-term in individuals originating from a wild population. This result is consistent with a previous correlational study where circulating baseline levels of IGF-1 were found to be positively associated with MDA in adult house sparrows [15]. Another study on nestling pied flycatchers found that daily IGF-1 injections increased the levels of the antioxidant enzyme glutathione peroxidase [16], which might either reflect lowered oxidative stress or up-regulated antioxidant activity in response to oxidative stress.

As IGF-1 concentration returned to pre-treatment levels at day 4, the difference in oxidative damage also disappeared between the groups. Microspheres were found to release encapsulated IGF-1 over several days in mice (e.g. [17]), whereas treatment effects disappeared by day 4 in our avian model, which indicates either a fast biodegradation of the microspheres or a strong negative feedback in reedlings (birds). This parallels findings of steady release hormone pellets that also have faster depletion in birds than in mammals [18].

The experimental increase in IGF-1 and MDA levels had no effect on long-term survival. This is probably due to the transient nature of the hormone peak. Although the experimentally elevated activity of the IIS pathway resulted in measurable increase in cellular oxidative damage, this short-term effect was probably too weak to affect survival on the long run. Remarkably, higher baseline IGF-1 (but not MDA) levels measured before the treatment were associated with lower mortality, not higher mortality as expected (see above); nonetheless, this association was weak and at the boundary of statistical significance. This result suggests that natural variation in IGF-1 levels may be the result of individual optimization (recently coined as the Optimal Endocrine Phenotype Hypothesis; [19]). In this context, high-quality individuals may afford to bear the costs of elevated IGF-1 levels (e.g. oxidative damage) while benefiting from its fitness-enhancing effects (e.g. boosting fecundity or being anti-inflammatory [20]).

We measured survival in a semi-natural environment under *ad libitum* diet regime and shelter from predators. Fluctuations in environmental conditions and stress stimuli may substantially reorganize the physiological network and, therefore, alter the adaptive value of a given endocrine phenotype [15]. IGF-1 levels showed high inter-individual variability and significant repeatability over three months indicating that the circulating levels of this hormone may be a consistent individual phenotypic marker. Whether individuals with naturally high IGF-1 levels also realize fitness advantages under more challenging natural conditions remains to be investigated.

## Ethics

The study was licensed by the local authorities.

## Data accessibility

All data supporting the results will be deposited at Dryad upon acceptance.

## Authors’ contributions

ÁZL and ZT conceived and conducted the experiment, ÁZL and ZT collected the samples and the data, ÁZL, ZT, JP and CIV measured the samples, SMK and BAG contributed reagents, ÁZL analysed the data, ÁZL and CIV wrote the article, all authors approved the final version.

